# DISTINCT NEURAL SIGNATURES OF HIPPOCAMPAL POPULATION DYNAMICS DURING LOCOMOTION-IN-PLACE

**DOI:** 10.1101/2025.10.28.685240

**Authors:** Samsoon Inayat, Brendan B. McAllister, Ian Q. Whishaw, Majid H. Mohajerani

**Affiliations:** Canadian Centre for Behavioural Neuroscience, University of Lethbridge, Lethbridge, Alberta, Canada; Department of Psychiatry, McGill University, Montreal, Quebec, Canada; Department of Psychology, University of Nevada, Las Vegas, USA; Department of Psychology, University of Calgary, Calgary, Alberta, Canada

## Abstract

Hippocampal CA1 neurons modulate their activity with movement variables such as time, distance, and speed, yet it remains unclear how these representations reorganize across behavioral states, from externally driven to self-paced movement and immobility. Here, we investigated how sensory events that initiate or terminate locomotion, structure CA1 population codes and how these codes reorganize across sensory-driven locomotion, spontaneous locomotion, and forced immobility. Using two-photon calcium imaging in head-fixed Thy1-GCaMP mice (n = 5) performing the air-induced running task on a non-motorized conveyor belt, we examined neuronal firing-rate modulation across a series of behavioral configurations designed to probe distinct forms of locomotion-in-place. In the No-Brake (locomotion-permitted) condition, the belt rotated freely, allowing animals to execute full cyclic limb movements in response to air stimulation. In the Brake (immobility) condition, the belt was fixed, restricting movement to partial or attempted locomotor motions. Firing-rate modulation with respect to time, distance, and speed was quantified using linear (Pearson correlation) and nonlinear (mutual information) metrics under permutation testing in the natural reference domains. Behaviorally, air stimuli produced faster, sustained running during air-on and more variable, self-paced movement during air-off. Neurally, a larger fraction of CA1 cells was active and significantly modulated during air-off. Within the modulated set, singularly tuned cells (time, distance, or speed) predominated over mixed-tuned cells, and speed-modulated cells peaked earlier after stimulus onset or offset than time- or distance-modulated cells. Under Brake, CA1 activity was predominantly singularly tuned to time or movement-in-place, with stronger movement modulation and engagement post-stimulation. Despite substantial single-cell turnover across configurations and phases, population-level analyses revealed a coherent, air-phase-locked organization and distinct movement-related populations across Brake and No-Brake conditions. These results indicate a state-dependent reweighting of sensorimotor features implemented atop a conserved ensemble scaffold in CA1.

## INTRODUCTION

Accumulating evidence indicates that ongoing sensorimotor events and movement modulate cellular activity in the hippocampal CA1 network^1–7^, providing a basis for its established roles in spatial behavior^8,9^, episodic learning and memory^10–14^, context representation^15–17^, and scene construction^18^. At a fundamental level, such modulation may arise through multiple mechanisms: (1) extrinsic or intrinsic influences, such as sensory events, environmental or internal to the body; (2) intrinsic cellular network dynamics; (3) spontaneous intracellular molecular processes; or (4) interactions among these mechanisms. Intrinsic network dynamics can themselves involve diverse processes, including inputs from other brain regions/networks carrying higher-order sensory features or motor efference copies, as well as computations within the hippocampal circuit that generate command-like outputs or predictive signals, or memory-related encoding, decoding, and indexing. Consistent with these mechanisms, a substantial body of research has shown that hippocampal neurons can fire transiently in relation to specific sensory or behavioral events ^19–26^, at fixed times^3,27–32^, fixed distances^27,29,31,33–36^, or in relation to spatial locations in two dimensional^37^, three-dimensional^38^, or even frequency-based environments^39^. In addition to spatial and temporal correlates, hippocampal neurons also encode task-relevant variables such as task type ^15,40,41^, behavioral parameters like speed ^42^, and internal cognitive states including motivation ^43^ and attention ^44^. To further elucidate the diversity of hippocampal cellular activity, it is essential to examine state-dependent, behavior-specific population dynamics both in the presence and absence of external sensory stimulation, an approach that remains underexplored and motivates the present study.

Beginning with the identification of differential features of local field potentials in the hippocampus ^4,5,45,46^ during immobility and locomotion, state-dependent cellular activity and encoding has been seen in both cortex ^47–53^ and hippocampus ^2,54–56^. Immobility is associated with large irregular activity and sharp-wave ripples, whereas locomotion is marked by theta oscillations, reflecting distinct network states. Correspondingly, hippocampal cells can fire at specific latencies from sensory events in both immobility ^30,57^ and locomotion ^3,29^ while locomotion additionally reveals distance-related firing from sensory or behavioral landmarks (e.g., reward sites) ^29,33–36^ and modulation by running speed^42,58,59^. These findings highlight that hippocampal coding principles are not fixed but reorganize across network states. During locomotion, however, time and distance are intrinsically confounded by speed. To disentangle these parameters, motorized treadmill studies have clamped either distance or time ^27,29,31,60^, revealing cells modulated by one, the other, or both. Yet, interpretation is complicated by vestibular input, treadmill-related proprioception, reward motivation, and initial acceleration, which can obscure genuine time or distance modulation. In natural behavior, animals alternate between immobility, stimulus-driven (e.g., evoked fleeing) and post-stimulus or self-paced exploration. Despite prior insights, it remains poorly understood how hippocampal representations of time, distance, speed, and movement in-place reorganize across immobility and locomotion states; whether speed dominates under externally imposed movement; whether external cues alter neuronal modulation; and whether single-cell identities and population-level organization persist or reconfigure across states.

In this study, we examine how hippocampal CA1 neuronal activity is modulated by time, distance, and speed in relation to sensory events that structure locomotor behavior, and we compare these representations with sensory-related time- and movement-modulated cellular activity during immobility. We build on the same dataset previously reported in *iScience*^2^ where we introduced the air-induced running (AIR) task in which head-fixed mice run on demand on a non-motorized conveyor belt in response to an air-stream applied to their back (Fig. 1A). After AIR training and initial testing, animals underwent seven behavioral configurations under Brake and No-Brake conditions while two-photon calcium imaging was used to record hippocampal CA1 activity (Fig. 1B, 1K-L). In the No-Brake condition, animals exhibited stimulus-driven locomotion; here we analyze cellular activity during alternating fixed-distance (air-on) and fixed-time (air-off) phases anchored to stimulus onset and offset, while minimizing reward influences and motorized-belt confounds. We further compare these event-related representations with cellular activity during Brake conditions to assess how hippocampal coding reorganizes across network states, providing new insight into how CA1 ensembles integrate sensory events with movement dynamics.

**Figure 1.**
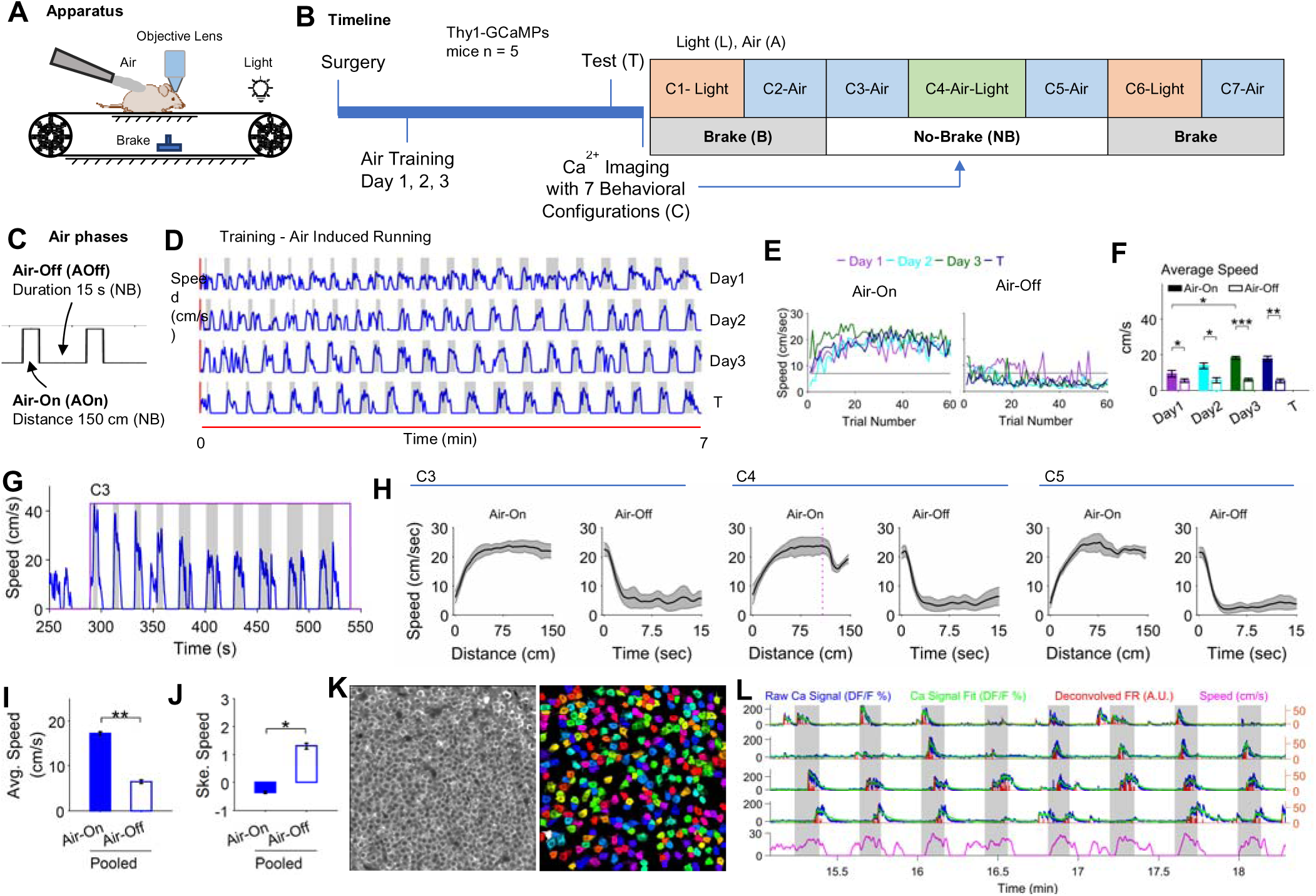
Air/Stimulus-induced running (AIR) task for studying hippocampal CA1 neuronal encoding. **(A)** Experimental setup. To motivate running, a head-fixed mouse on a non-motorized linear conveyor belt receives a mild air stream on its back. **(B)** Experiment timeline; surgery (CA1 cranial window), recovery (∼3 weeks), training (∼30 mins for 3 days), behavioral test ∼2 weeks after training (30 mins), calcium imaging session next day with mice experiencing 7 configurations (Brake - B and No-Brake -NB conditions). **(C)** Air phase structure; for the NB condition, mice ran a fixed distance (150cm) in response to air application and waited for a fixed time interval of 15s in between air applications. For the B-condition, air is ON and OFF for 5 s and 10 s respectively **(D)** Raw traces of the speed from a representative animal on training days (first 7 mins shown). Gray shaded regions indicate trials of air stream application and white regions indicate air-off phases. **(E)** Average speed during trials vs trial numbers for both air-on and air-off phases for training and testing day for a representative animal. **(F)** RM-ANOVA result for average speed versus training days during air-on and air-off phases. **(G)** Representative trace from an animal showing speed vs time for Configurations C3. **(H)** The average speed during air-on and air-off for C3, C4, and C5 averaged over trials and animals. Thick line shows average speed while shaded region shows SEM. The dotted red line shows onset of 200 ms light flash at 110 cm in C4 during the air-on phase. **(I-J)** RM-ANOVA result of the average and skewness of speed respectively during air phases across C3, C4, and C5. **(K)** Average image (left) of the calcium imaging time series from a representative animal, and arbitrarily color-coded regions of interest indicating identified cell bodies (right) in a ∼400 × 400 μm imaging window (from Suite2P). **(L)** Calcium signal (raw and fitted), deconvolved firing rate (FR), and speed vs time for representative cells. Shaded regions show air-on phase. Error bars = SEM. n = 5 mice, *p < .05, **p < .01, ***p < .001

## RESULTS

### Air events structure locomotion behavior in the air-induced running (AIR) task

As reported previously, an air-induced running task^2^ was developed to enable quick learning of head-fixed mice to run on a non-motorized treadmill in response to an air stream applied to their back (Fig. 1A). Mice trained for three days and tested once ran in response to air application during the air-on phase, continued to move briefly post-stimulation and spontaneously locomoted during the air-off phase (Fig. 1C, D, E). A repeated-measures analysis of variance (RM-ANOVA) examining the effects of Day × Air Phase (4 x 2) on average trial speed revealed a significant interaction [F(3,12) = 25.33, p < 0.001, η² = .30]. Posthoc analyses with Tukey’s honestly significant difference (HSD) confirmed that while speed increased over days during the air-on phase, it remained stable in the air-off phase (Fig. 1F) suggesting that mice learned to modulate their speed in response to the presence or absence of air.

After training and initial testing, animals experienced seven behavioral configurations in Brake (B) and No-Brake (NB) conditions (Fig. 1B) while calcium imaging was performed. In each of the Configurations 3, 4, and 5 in NB condition, animals completed 10 trials similar to those on the testing day (Fig. 1C), in which they ran a fixed 150 cm lap of the treadmill during the air-on phase, followed by a 15 s air-off phase before the next trial (Fig. 1G and 1H). In Configuration 4, a 200 ms light flash was also presented 110 cm from the air onset. An RM-ANOVA examined the effect of Configuration × Air Phase × Trial (3 x 2 x 10) on average trial speed and a significant main effect of Air Phase was observed [F(1,4) = 46.94, p = .002, η² = .58] with posthoc analyses showing higher speeds during the air-on phase compared to the air-off phase (Fig. 1I). A similar analysis for the skewness of trial speed showed a main effect of Configuration on skewness [F(2,8) = 9.02, p = .009, η² = .03] but no pairwise differences and a main effect of Air Phase [F(1,4) = 14.17, p = .020, η² = .49], with negative and positive skewness during the air-on and air-off phases respectively (Fig. 1J). The latency of movement after air-onset (with speed > 0 cm/s) during the air-on phase was 0.470 ± 0.059 s (mean ± SEM) and was not significantly different across configurations and trials (RM-ANOVA, no main or interaction effects). The latency of movement after air-offset during the air-off phase was 0 s for all trials and configurations indicating that the animals were moving at the time of air-offset. During the air-off phase, time taken to reach immobility state (0 cm/s speed) after air-offset was (mean ± SEM: 5.209 ± 0.298) and was not significant across configurations and trials. The time to complete the air-on phase (mean ± SEM: 8.98 ± 0.24 s) was significantly smaller than the fixed 15 s duration of the air-off phase, [F(1,4) = 33.39, p = .004, η2 = .69]. The distance covered during the air-off phase (mean ± SEM: 87.89 ± 6.05 cm) was not significantly different from the fixed 150 cm distance covered during the air-on phase (RM-ANOVA, no main or interaction effects).

These results suggest that the air-on and air-off phases represent distinct movement patterns, with animals exhibiting a structured, stimulus-driven response during the air-on phase, continued locomotion after air-offset, and more variable, spontaneous movement during rest of the air-off phase. The negative skewness observed in the air-on phase suggests that mice maintained higher speeds for longer durations, whereas the positive skewness in the air-off phase reflects a bias toward lower speeds, consistent with a more exploratory or resting state.

### Larger population activity in the post-stimulation air-off phase with larger population of singularly modulated cells firing in relation to time, distance, and speed

To investigate how hippocampal CA1 activity relates to movement variables, we examined the modulation of cellular firing rates (FR) in relation to time, distance, and speed during the air-on and air-off phases of Configurations 3, 4, and 5 in the No-Brake condition (Fig. 1B). FR was estimated by deconvolving calcium fluorescence signals (Fig. 1L) and was binned in both time and distance (Fig. 2). Time binning was used to assess time and speed modulation, whereas distance binning was used to assess distance modulation so that each variable was analyzed in its natural reference domain. For visualization, plots of the fixed-distance air-on and fixed-time air-off phases are displayed with distance and time binning, respectively, for convenience.

**Figure 2.**
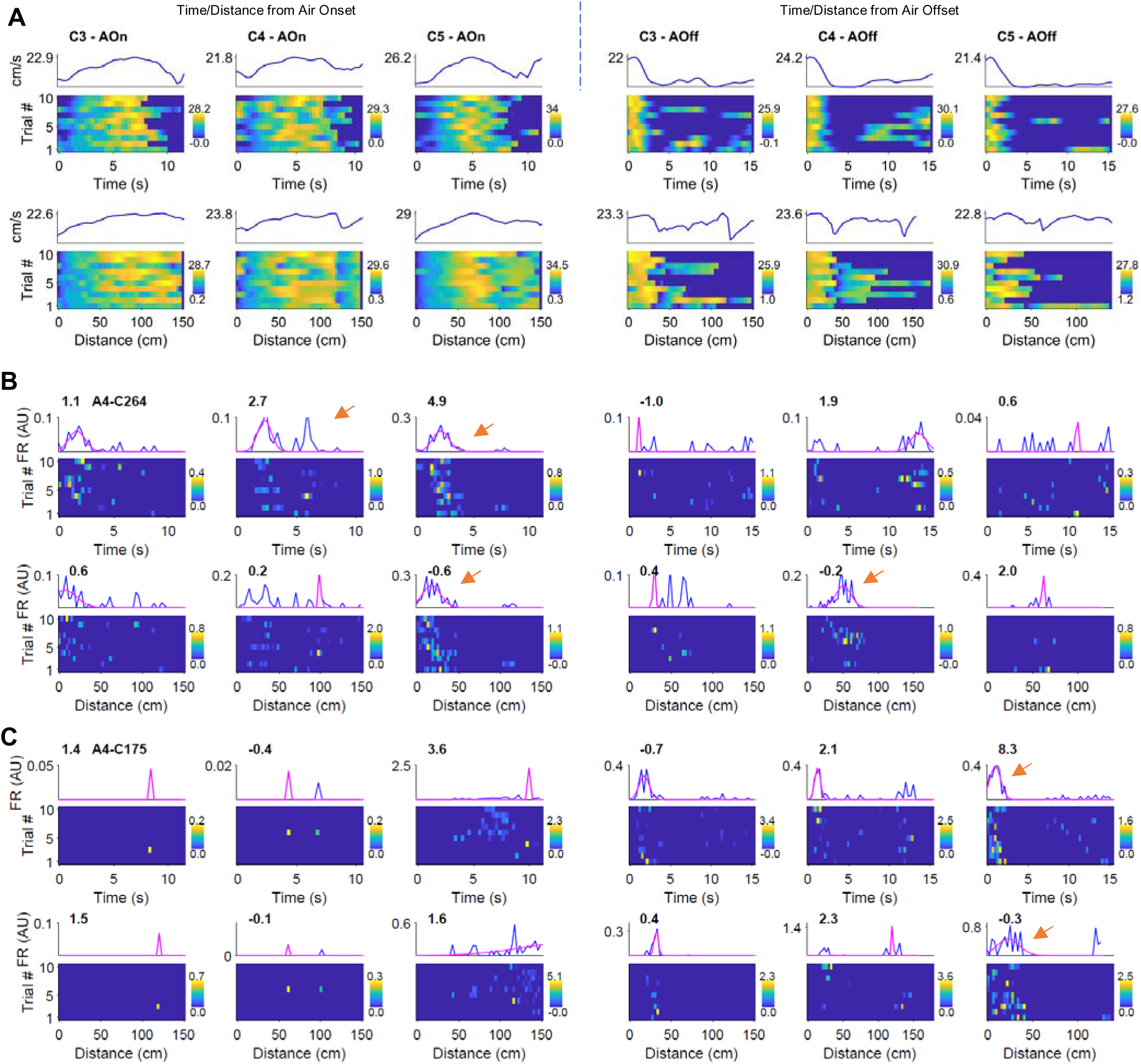
Representation of speed and neuronal firing rate in relation to sensory events air-onset and air-offset in the air-on and air-off phases respectively with both time- and distance-binning. **(A)** Raster plots of speed (bottom) and average speed (top) for a representative animal for configurations C3, C4, and C5 and for both time and distance binned data sets. **(B)** Bottom panels show raster plots of firing of a representative cell (corresponding to panels in A for the same animal). In the top plots, blue lines show means over trials. Magenta lines show 1D Gaussian fit on the mean response. Time and/or distance modulation of firing rate can be seen (red arrows). **(C)** same as in B for another representative cell. The numbers above the graphs represent Z-scored mutual information (zMI).

The activity dynamics of the entire cellular population, visualized through rate vector maps, revealed more transient activity during the air-on phase compared to the air-off phase (Fig. 3A, top row, representative animal). Population vector correlation plots showed a tighter diagonal “band” during air-on, indicating stronger correlations among adjacent bins consistent with short-lived, transient firing patterns (Fig. 3A, middle row; averaged across animals, bottom row). Because the full population included sparsely active cells (Fig. 3A), we focused subsequent analyses on cells that fired in at least three air trials (response fidelity > 30%; Fig. 3B). These “good RF” cells were used to ensure reliable assessment of FR modulation by movement variables. A two-way RM-ANOVA (Configurations × Air-Phase; 3 × 2) on the percentage of good RF cells revealed a significant main effect of Air-Phase [F(1,4) = 9.54, p = 0.037, η² = 0.49], with a larger fraction of active cells in the air-off phase (67.36 ± 1.72 %) than in the air-on phase (49.22 ± 3.16 %).

**Figure 3.**
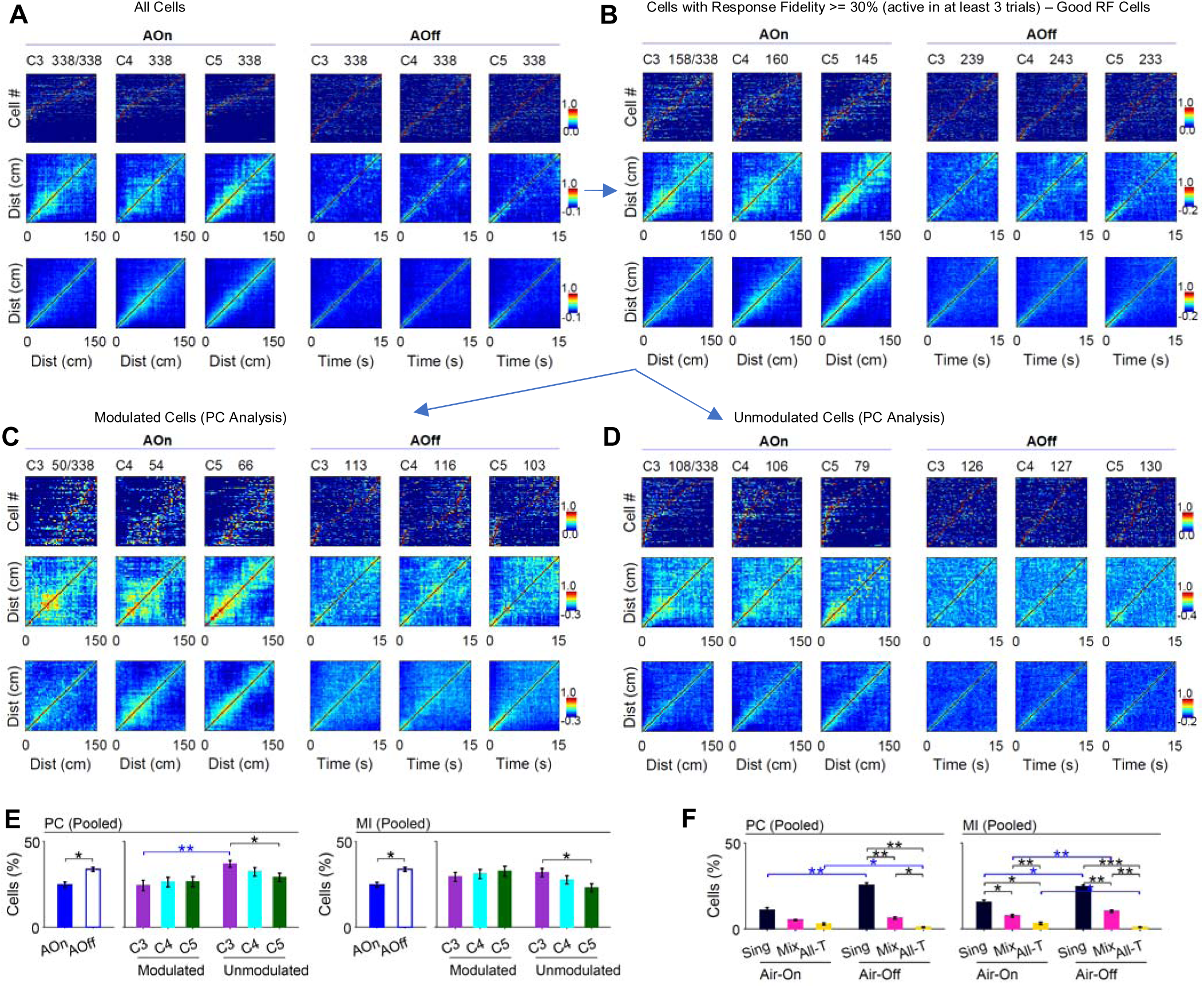
Cellular population plots and identification of time-, distance-, and speed-modulated cells. **(A)** All recorded cells from a representative animal. Rate-vector maps (cells sorted by peak) shown in the top row. Population-vector correlation matrices (middle row) corresponding to panel A (brighter diagonal = more transient/structured activity). Average population-vector correlation (n = 5 mice), bottom row **(B)** Same as panel A but for Good-RF cells (response fidelity ≥30%, active in ≥3 trials). **(C)** Same as panel A but for modulated cells (permutation-tested) by time, distance, or speed using PC metric. **(D)** Same as panel A but for unmodulated cells from the same sessions. (E–J) Group summaries (n=5 animals). **(E)** RM-ANOVA result comparing the percentage of modulated vs. unmodulated cells across configurations and air-phases. **(F)** RM-ANOVA comparing percentage of singularly-(Sing.), mixed-(Mix.), and all-modulated (All-T) cells. Bars show mean ± SEM; HSD post-hoc comparisons. AOn/AOff, air-on/air-off; PC, Pearson correlation; MI, mutual information. n = 5 mice, *p < .05, **p < .01, ***p < .001

We next quantified how these firing rates related to behavioral/movement variables of time, distance, and speed. Linear relationships between FR and each variable were quantified using Pearson correlation (PC), and non-linear relationships using mutual information (MI). Significance was determined by permutation testing, in which observed PC or MI values were compared to null distributions generated by 1,000 random shuffles of FR; a cell was considered significantly modulated if its metric exceeded the top 5 % of the null (p < 0.05; zPC or zMI > 1.645, one-tailed).

Three-way RM-ANOVAs (Modulation × Configuration × Air-Phase; 2 × 3 × 2) for percentage of modulated vs unmodulated cells were performed separately for PC and MI revealing a consistent pattern. For both metrics, there was a significant main effect of Air-Phase [PC: F(1,4) = 9.54, p = 0.037, η² = 0.35; MI: F(1,4) = 9.54, p = 0.037, η² = 0.34], indicating that a larger proportion of CA1 cells were active and modulated in the post-stimulation air-off phase compared to the air-on phase (Fig. 3E).

Additionally, a significant Modulation × Configuration interaction was observed [PC: F(2,8) = 7.04, p = 0.044, η² = 0.10; MI: F(2,8) = 5.84, p = 0.037, η² = 0.14]. Post hoc analyses revealed a gradual decline in unmodulated cells and a trend toward increasing modulated-cell fractions across configurations, although the latter did not reach significance (Fig. 3E).

To further characterize the structure of modulation in hippocampal CA1, we classified modulated neurons based on the number of behavioral variables (time, distance, speed) to which their firing was significantly related. Cells that were significantly modulated by only one variable were labeled singularly modulated, those modulated by two variables were labeled mixed-modulated, and those modulated by all three variables were labeled all-modulated. The percentage of these cells was then compared using a three-way RM-ANOVA; Modulation-Order × Configuration × Air-Phase (3 × 3 × 2) separately for the PC and MI metrics. There was a strong Modulation-Order × Air-Phase interaction (PC: [F(2,8) = 49.44, p < 0.001, η² = 0.51]; MI: [F(2,8) = 19.33, p < 0.001, η² = 0.35]) with post hoc comparisons revealing that singularly modulated cells constituted the largest population in both phases and increased significantly in proportion during the post-stimulation air-off phase (Fig. 3F). In contrast, mixed-modulated cells showed little or no phase-related change, and all-modulated cells represented only a small fraction of the overall population. The pattern was consistent across both linear (PC) and non-linear (MI) metrics, confirming that hippocampal CA1 activity becomes more selectively tuned after stimulus offset.

Together, these findings indicate that a larger CA1 population is engaged in the air-off phase, when movement continues after stimulus offset, and that experience across configurations is associated with gradual reorganization of modulated cells. Within the modulated population, there is a larger percentage of singularly modulated and a larger presence in the air-off phase. The shift toward single-variable modulation suggests that post-stimulation network states enhance feature-specific encoding of time, distance, and speed within the CA1 ensemble. In the subsequent analysis, only singularly modulated cells are further analyzed.

### Earlier peak firing of speed modulated cells after stimulus events compared to time and distance modulated cells

The distribution and characteristics of time, distance, and speed modulated cells in the No-Brake condition (Configurations 3, 4, and 5) was investigated by examining their percentages and firing properties. Rate vector maps and population correlation plots for time-, distance-, and speed-modulated cells showed varied populations active across configurations and air-phases with variable characteristics including both transient and sustained firing (Fig. 4A-F). Time- and distance-modulated cells in contrast with speed-modulated cells, were not reliably detected with the PC metric (see average population correlation plots). As noted, cells fired more transiently (tighter glow around diagonals) during air-on compared to during air-off phase but detection was overall better for air-off phase. The population of cells also seemed to differ in the location of peak firing. To quantitatively compare these populations, the response variables; percentages of cells and location of peak firing were compared using RM-ANOVAs.

**Figure 4.**
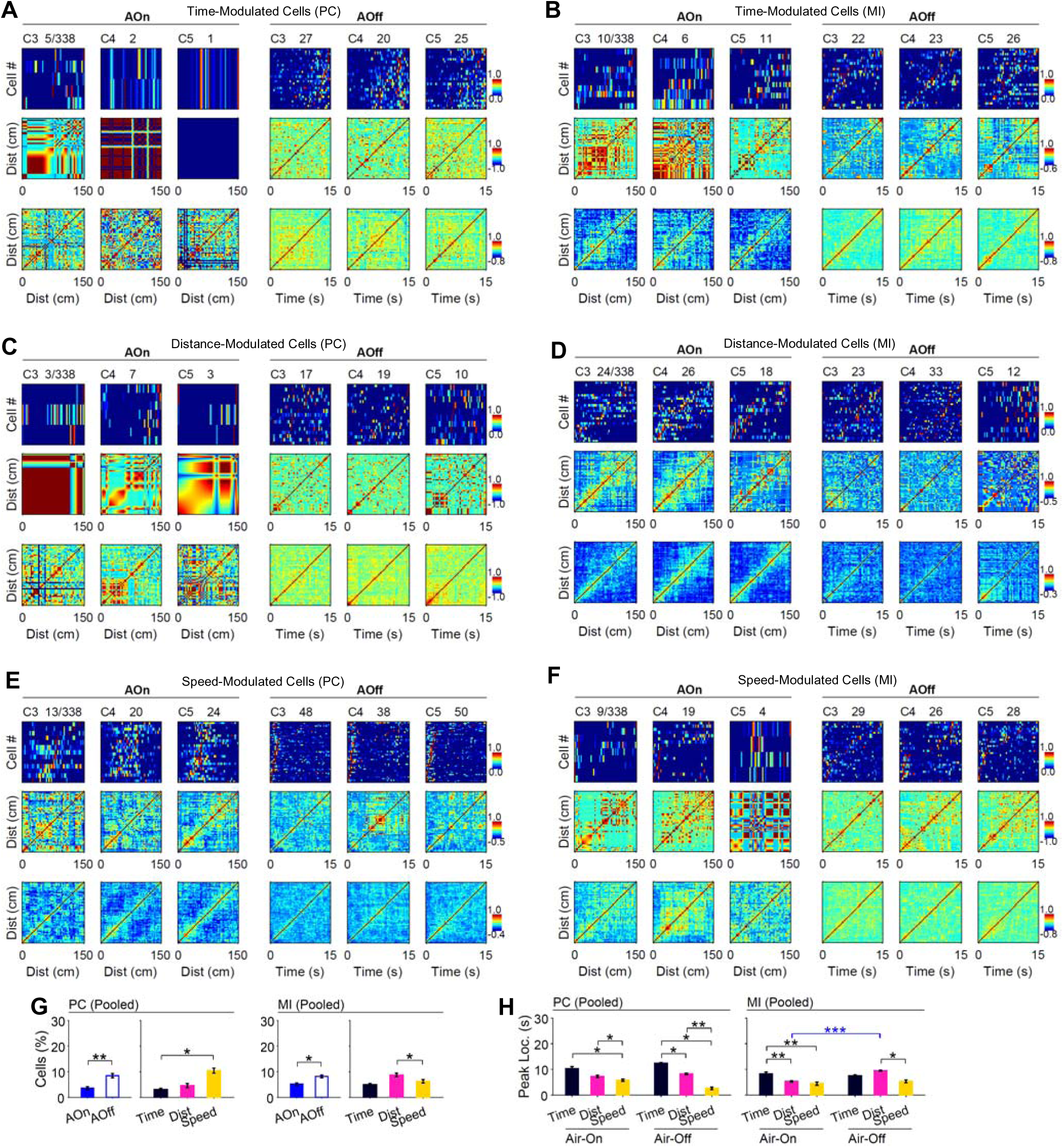
Cellular population plots of representative time-, distance-, and speed-modulated cells. **(A)** Time-modulated cells identified with the PC metric. Rate-vector maps (cells sorted by peak) shown in the top row. Population-vector correlation matrices (middle row) corresponding to panel A. Average population-vector correlation (n = 5 mice), bottom row **(B)** Same as panel A but for time-modulated cells identified with the MI metric. **(C-D)** Same as panels A-B but for distance-modulated cells. **(E-F)** Same as panel A-B but for speed-modulated cells. **(G)** RM-ANOVA result comparing the percentage of modulated cells across configurations and air-phases and type of modulation (time, distance, or speed). **(H)** RM-ANOVA result comparing the peak location of firing in seconds from air-onset or air-offset across type of modulation (time, distance, or speed) and across configurations and air-phases. Bars show mean ± SEM; HSD post-hoc comparisons. AOn/AOff, air-on/air-off; PC, Pearson correlation; MI, mutual information. n = 5 mice, *p < .05, **p < .01, ***p < .001

For the percentage of cells, a three-way RM-ANOVA (Modulation-Type × Configuration × Air-Phase; 3 × 3 × 2) separately for PC and MI metrics revealed main effect of Modulation-Type (PC: [F(2,8) = 9.22, p = .008, η^2^ = .47]; MI: [F(2,8) = 7.84, p = .013, η^2^ = .23]). Post hoc comparisons revealed that with the PC metric, speed-modulated cells outnumbered time-modulated cells, whereas with the MI metric, distance-modulated cells exceeded speed-modulated cells. (Fig. 5A). There was also a main effect of Air-Phase (PC: [F(1,4) = 72.46, p = .001, η^2^ = .34]; MI: [F(1,4) = 16.04, p = .016, η^2^= .22]) and posthoc comparisons revealed a larger percentage of cells in the air-off phase (Fig. 4G).

**Figure 5.**
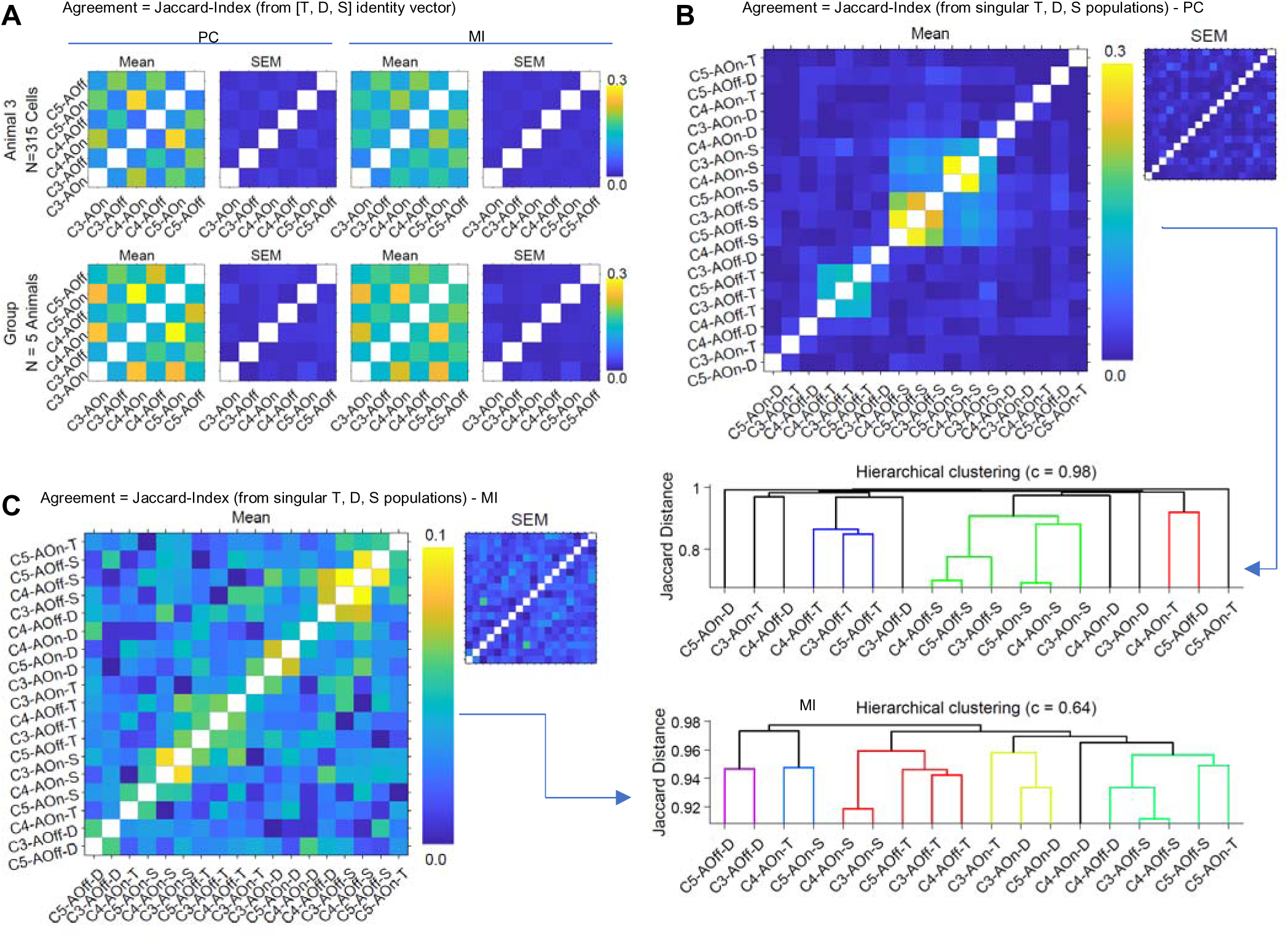
High single-cell modulation turnover but coherent population-level organization across configurations and air phases. **(A)** Agreement of cell-modulation identities across configurations and air-phases. For each cell, a 3-element modulation-identity vector [T,D,S] was compared across the six cases using the Jaccard index (0 - 1). Heat maps show mean and SEM for a representative animal (top) and the group (bottom, n=5 mice). Brighter AOn–AOn and AOff–AOff blocks indicate higher within-phase agreement than across-phase (AOn–AOff). **(B)** Population-level agreement for cells identified with the PC metric. Pairwise Jaccard agreement among the 18 singularly-modulated populations (3 labels × 3 configurations × 2 phases). Heat map shows mean agreement (inset: SEM); hierarchical clustering on 1–agreement (Jaccard distance) yields modulation/phase groupings (dendrogram; cophenetic correlation c=0.98). **(C)** same as panel B but for cells identified with the MI metric. Group data, n = 5 mice.

Peak locations of firing were determined in time-binned data set as time of peak firing from sensory events (air onset or offset) in the mean response and were compared using three-way RM-ANOVAs (Modulation-Type x Configuration x Air-Phase; 3 × 3 × 2) separately for PC and MI metrics. There was a significant Modulation-Type × Air-Phase interaction (PC: [F(2,4) = 63.50, p < 0.001, η^2^ = .42]; MI: [F(2,8) = 8.45, p = .011, η^2^ = .26]). Posthoc comparisons showed significantly smaller values of peak locations for the speed modulated cells compared to time and distance modulated cells for both air-on and air-off phases (Fig. 4H).

Together, these findings reveal a structured organization of modulation within dorsal CA1, where speed-modulated cells form a dominant and temporally earlier component of the population response relative to time- and distance-modulated cells. No effect of configuration was observed in the differential activation of time-, distance-, or speed-modulated cells.

### High single-cell modulation turnover but coherent population-level organization across configurations and air phases

To investigate the dynamics of modulation by time-, distance-, and speed-modulated cells across Configurations 3, 4, and 5 (No-Brake condition) and air phases, we computed agreement (= Jaccard Index) between cellular and population identity vectors (see methods). The Jaccard-index quantifies overlap between two identity vectors (0 = no overlap; 1 = full overlap). For each cell, an identity vector with three elements [T, D, S] corresponding to time-, distance-, and speed-modulation was defined.

Identity vectors were compared across the six cases (3 configurations × 2 air phases) and the resulting agreement values were averaged over cells and then over animals to summarize identity dynamics (Fig. 5A). Agreement was generally modest (< 0.3) consistent with high single-cell turnover (cells often changed modulation identity across cases). Nevertheless, the average agreement matrices showed a structured checkerboard pattern with higher values for same-phase pairs (AOn-AOn and AOff-AOff) than cross-phase pairs (AOn-AOff). Quantifying this contrast, same-phase minus cross-phase agreement (Δ) was positive in every animal for both metrics (PC: Δ = 0.122 ± 0.022 SEM, Wilcoxon signed-rank one-sided p = 0.031; MI: Δ = 0.078 ± 0.009 SEM, p = 0.031), indicating that identities are more similar within phase than across phases. Thus, despite low absolute agreement reflecting frequent identity changes, the population exhibits a coherent, phase-dependent organization across configurations.

Next, we compared population identities for time-, distance- and speed-modulated cells across configurations and air-phases by computing the Jaccard-index between singularly modulated populations (exactly one of T/D/S) in each case, yielding an 18 x 18 agreement matrix (3 cell-type labels x 3 configurations x 2 air-phases). For PC, the matrix showed modest absolute values (< 0.3) consistent with turnover, but a second-order hierarchical-clustering analysis on 1-agreement (Jaccard distance) revealed distinct groups (e.g., speed-related and time-related clusters; Fig. 5B). The group dendrogram fit was high (cophenetic correlation c = 0.98). For MI, absolute agreements were smaller (color scale 0 – 0.1), yet the same analysis still revealed structured grouping of populations (Fig. 5C) with a moderate tree fit (c = 0.64).

Together, these results suggest that although overall single-cell modulation shows substantial turnover across configurations and air-phases, there is a coherent population-level organization. Cells tend to retain identity within the same air phase (AOn ➔ AOn, AOff ➔ AOff) more than across phases (AOn ➔ AOff), and second-order analyses indicate higher-order structure across labels, with the PC (linear) analysis showing a stronger organization than MI (non-linear).

### Larger population of singularly modulated cells in relation to time and movement in-place in the Brake Condition under forced immobility

To examine the modulation of neural activity in the air-on and air-off phases in the Brake Condition (Configurations 2 and 7), time- and movement-modulated cells were identified using the permutation testing with PC and MI metrics as described above (Fig. 6 A-D). Movement-modulated cells in the Brake condition were identified using the speed signal of the conveyor belt which although was clamped was still displaced back and forth slightly in-place by small animal paw movements perturbing the rotary encoders.

**Figure 6.**
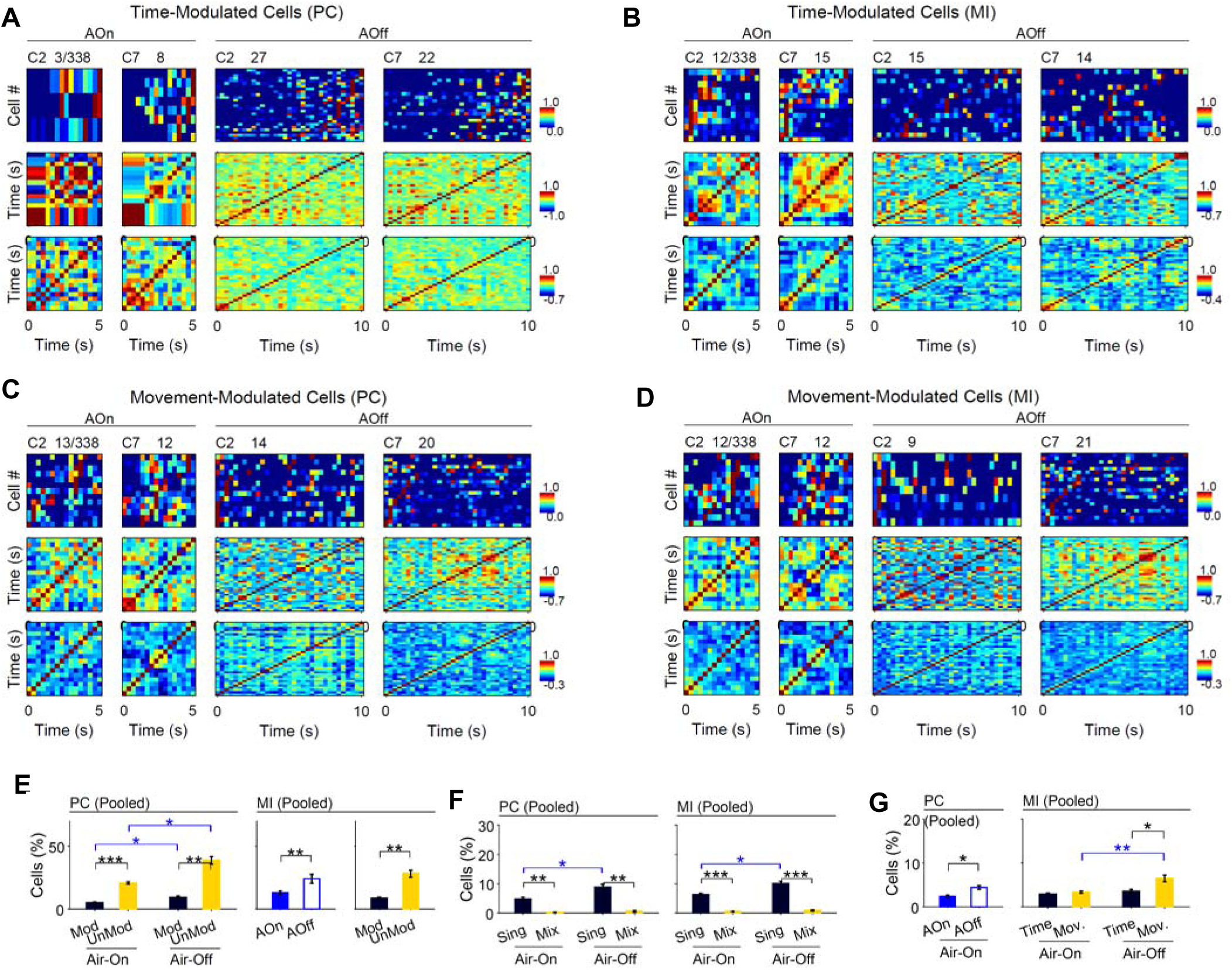
Cellular population plots of representative time- and movement-modulated cells. **(A)** Time-modulated cells identified with the PC metric. Rate-vector maps (cells sorted by peak) shown in the top row. Population-vector correlation matrices (middle row) corresponding to panel A. Average population-vector correlation (n = 5 mice), bottom row **(B)** Same as panel A but for time-modulated cells identified with the MI metric. **(C-D)** Same as panels A-B but for movement-modulated cells. **(E)** RM-ANOVA result comparing the percentage of modulated vs. unmodulated cells across configurations and air-phases. **(F)** RM-ANOVA comparing percentage of singularly-(Sing.) and mixed-(Mix.) cells across configurations and air-phases. **(G)** RM-ANOVA result comparing the percentage of modulated cells across configurations and air-phases and type of modulation (time or movement). Bars show mean ± SEM; HSD post-hoc comparisons. PC, Pearson correlation; MI, mutual information. n = 5 mice, *p < .05, **p < .01, ***p < .001

The percentage of modulated and unmodulated cells was compared with three-way RM-ANOVAs (Modulation × Configuration × Air-Phase; 2 × 2 × 2) separately for PC and MI metrics. For the PC metric, there was a significant Modulation × Air-Phase interaction [F(1,4) = 10.06, p = .034, η2 = .32] whereas for the MI metric, there were main effects; Modulation [F(1,4) = 48.58, p = .002, η2 = .73] and Air-Phase [F(1,4) = 24.14, p = .008, η2 = .48]. Posthoc comparisons showed a larger percentage of unmodulated cells overall and larger percentage of modulated cells in the post-stimulation air-off phase compared to the air-on phase (Fig. 6E).

Next, the percentages of singularly- and mixed-modulated cells, as for the No-Brake conditions (see above), were compared with three-way RM-ANOVAs (Modulation-Order × Configuration × Air-Phase; 2 × 2 × 2). For both the PC and MI metrics, there was a significant Modulation-Order × Air-Phase interaction (PC: [F(1,4) = 14.46, p = .019, η2 = .23]; MI: [F(1,4) = 15.63, p = .017, η2 = .30]). Posthoc comparisons revealed a larger percentage of singularly-tuned cells and a larger presence in the air-off phase (Fig 6F).

Time- and movement-modulated cells were then compared with three-way RM-ANOVAs separately for PC and MI metrics; Modulation-Type × Configuration × Air-Phase (2 × 2 × 2). A larger percentage of modulated cells in the air-off phase was found; main effect of Air-Phase (PC; [F(1,4) = 19.21, p = .012, η^2^ = .32], MI; [F(1,4) = 17.70, p = .014, η^2^ = .36]) and Modulation-Type × Air-Phase interaction effect for MI only [F(1,4) = 14.66, p = .019, η2 = .19] (see posthoc comparisons; Fig. xx). Furthermore, in the air-off phase, the percentage of movement-modulated cells was larger than time-modulated cells (Fig. 6G). These results suggest that the modulation specificity of neural activity in a given case e.g., configuration and air-phase, was largely singular i.e., either time- or movement-modulated. Overall, there was a larger activation during the air-off phase as was also observed for the No-Brake conditions.

### Coherent phase-dependent population-level organization within Brake-condition distinct from organization in the No-Brake condition

The population-level organization of time- and movement-modulated cells in the Brake condition was first investigated with an identity vector with two elements [T, M] corresponding to time- and movement-modulation (as was done in the No-Brake condition). Identity vectors were compared across four cases (two configurations × two air phases: C2–AOn, C2–AOff, C7–AOn, C7–AOff), and the resulting agreement (= Jaccard Index) values were averaged first across cells and then across animals to summarize modulation identity dynamics (Fig. 7A). Agreement values were modest overall (< 0.5), consistent with frequent changes in modulation identity across conditions. Nonetheless, both single-animal and group matrices displayed a structured pattern, with stronger within-phase (AOn–AOn, AOff– AOff) than cross-phase (AOn–AOff) similarity. Quantifying this contrast, the same-phase minus cross-phase difference (Δ) was positive in all animals for PC (mean ± SEM = 0.065 ± 0.022; Wilcoxon signed-rank one-sided p = 0.031) and showed a similar but weaker trend for MI (0.054 ± 0.027; p = 0.063). These results indicate that, even under restricted movement, CA1 populations preserve a coherent phase-dependent organization, with cell-specific modulation identities remaining more consistent within a behavioral phase than across phases.

**Figure 7.**
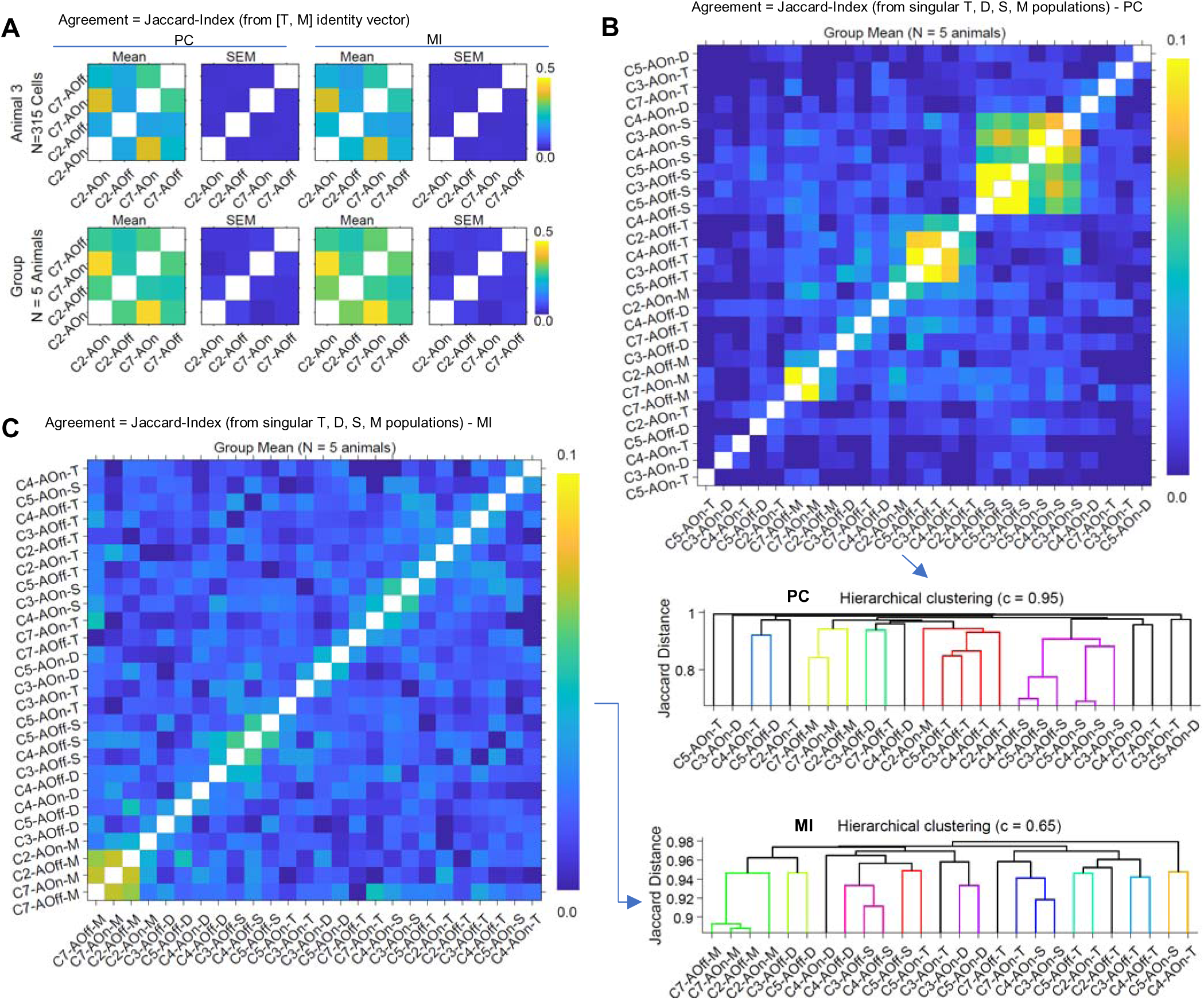
Coherent population-level organization across configurations and air phases in the Brake condition with distinct organization from No-Brake condition. **(A)** Agreement of cell-modulation identities across configurations and air-phases. For each cell, a 3-element modulation-identity vector [T,M] was compared across the four cases using the Jaccard index (0 - 1). Heat maps show mean and SEM for a representative animal (top) and the group (bottom, n=5 mice). Brighter AOn–AOn and AOff–AOff blocks indicate higher within-phase agreement than across-phase (AOn–AOff). **(B)** Population-level agreement for cells identified with the PC metric. Pairwise Jaccard agreement among the 26 singularly-modulated populations (3 labels × 3 configurations × 2 phases + 2 labels × 2 configurations × 2 phases). Heat map shows mean agreement (inset: SEM); hierarchical clustering on 1–agreement (Jaccard distance) yields Condition/modulation/phase groupings (dendrogram; cophenetic correlation c=0.95). **(C)** same as panel B but for cells identified with the MI metric. Group data, n = 5 mice.

Populations of time- and movement-modulated cells in the Brake condition were then compared with populations of time-, distance-, and speed-modulated cells in the No-Brake condition using Jaccard-index based analysis as described above for only No-Brake condition (Fig. 5B-C). Although there was a high-turnover for the identity of cells as assessed from low Jaccard-index values but the secondary clustering analysis revealed a coherent population-level organization across Brake and No-Brake conditions with distinct representations of movement-modulated cells (Fig. 7B and 7C, see clusters for S and M related cellular populations).

Taken together, these results suggest that within the individual Brake and No-Brake conditions, there is a coherent population-level organization of cellular responses across the air-phases but when compared across the two conditions, there is a distinct representation of movement-modulated cells.

## DISCUSSION

We investigated sensory event- and movement-related modulation of hippocampal CA1 neuronal activity across sensory-driven locomotion, spontaneous locomotion, and forced immobility. Using a head-fixed, cue-minimal paradigm that combined the air-induced running (AIR) task with two-photon calcium imaging, we examined how CA1 neurons represent movement-related variables (time, distance, and speed) within and across these behavioral states. In the No-Brake (locomotion-permitted) condition, animals alternated between a sensory-driven air-on phase, characterized by externally evoked running, and a post-stimulation air-off phase, marked by post-stimulation continued and self-paced spontaneous movement. The air stimulus organized motion with higher speeds during the air-on phase, however, larger neuronal populations were active during the air-off phase with singularly modulated cells dominating. Neural analyses revealed speed-dominant coding near sensory-events that organized locomotion, with time/distance-dominant coding farther from these events. In the Brake (immobility) condition, animals remained forcibly stationary yet displayed subtle in-place paw movements accompanied by dominant movement-related neural modulation. Despite substantial single-cell turnover across air-phases and sensory and behavioral states, the CA1 ensemble maintained a coherent, state- and phase-dependent organization, particularly related to movement, suggesting that dynamic reweighting of sensorimotor features occurs on top of a stable population scaffold.

In our AIR task, air stimulus functioned as an external drive that organized locomotion with higher speeds during air-on in the No-Brake (locomotion permitted) condition. Mice adopted a simple “run-while-air-present” policy and maintain near-stead speeds after an initial rise. Even though speeds were larger during air-on, a larger population of cells was active during the air-off phase. This may be due to the longer time mice spent in the air-off phase and is consistent with previous studies reporting that spontaneous and exploratory locomotion enhances CA1 ensemble participation ^32,61^. A larger cellular population was also seen in the air-off phase in the immobility condition suggesting a similar conclusion that longer post-stimulation time engages more cells.

Our study further supports the idea that neuronal representations persist across sensory-states (air-on vs air-off) as well as across behavioral states (immobility vs locomotion). Distinct neuronal representations across behavioral states are consistent with earlier work on hippocampal state dependence which described transitions between immobility-associated sharp-wave ripple states ^45,62^ and locomotion-associated theta oscillations ^4,5^. Furthermore, this population-level analyses agrees with prior observations that hippocampal ensembles maintain stable topologies even as constituent neurons remap ^15,32,55^ and that hippocampal representations depict multiple actions^6,7^. These findings suggest that hippocampal coding stability arises not from fixed cell membership but from preserved relational patterns among subpopulations.

The dominance and earlier peak firing of speed-modulated cells relative to time- and distance-modulated populations indicate that locomotor signals may provide the temporal scaffold upon which other variables are encoded. Previous studies have described speed as both a gain factor ^42,63^ and as a content variable influencing hippocampal rate and phase coding ^29,36^. Our data refine this view by suggesting hierarchical recruitment process: speed cells activate first, potentially informing downstream time and distance integrators as behavior evolves. This supports the notion that hippocampal representations of time and space may emerge from recursive transformations of self-motion signals ^34,62^.

The larger movement-in-place-compared to time-modulation in the forced immobility condition supports the idea that the hippocampal network contributes to sensorimotor integration, even in the absence of overt locomotion. Such modulation resembles hippocampal activity during consummatory behaviors, when animals exhibit small repetitive movements (e.g., licking, chewing, grooming) accompanied by sharp-wave ripples ^45,55^.

Our AIR paradigm characterized by cue-minimal, event-locked, and non-motorized design offers advantages over conventional treadmill paradigms by eliminating confounds from reward, vestibular input, and imposed kinematics ^2,28,29,60,64^. However, head fixation inherently limits vestibular and naturalistic feedback, which may constrain the full expression of sensorimotor and spatiotemporal coding. Furthermore, neural coding could partly arise from residual sensory cues, such as urine and feces, although these effects were minimized by cleaning and drying belts before each session. Future iterations incorporating fixed-time air-on phases or freely moving adaptations could better dissociate intrinsic time from sensory-driven distance encoding. Additionally, since calcium imaging underrepresents fast spiking events and subthreshold dynamics, combining imaging with simultaneous LFP recordings could clarify the relationship between cellular modulation and network oscillatory states.

Together, our findings highlight a state-dependent reweighting of sensorimotor features in the hippocampal CA1 with speed predominating under externally driven locomotion near sensory events, whereas time and distance dominating self-paced movement farther from sensory events. Despite single-cell reorganization, the ensemble retains coherent, phase-locked structure, suggesting that dynamic representations are implemented on top of a stable coding scaffold. This is also established in the immobile state with further finding of movement-in-place related neural representations distinct from movement-modulated neural activity during locomotion.

## RESOURCE AVAILABILITY

### Lead contact

Further information and requests for resources and reagents should be directed to and will be fulfilled by the lead contact, Majid H. Mohajerani. Materials availability: This study did not generate new unique reagents.

### Data and code availability

- This study did not generate any standardized datatypes. However, calcium imaging data reported in this paper will be shared by the lead contact upon request.
- All codes used for analyzing data in this paper will be made available upon the acceptance of the paper

## EXPERIMENTAL MODEL AND SUBJECT DETAILS

All animal procedures were approved by the Animal Welfare Committee of the University of Lethbridge and followed Canadian Council on Animal Care guidelines. Data were obtained from n = 5 adult (5–8 months old) Thy1-GCaMP6s [C57BL/6J-Tg(Thy1-GCaMP6s)GP4.3Dkim/J] mice (3 male, 2 female), as described previously^2^. This transgenic line expresses GCaMP6s in excitatory neurons across cortical and subcortical regions. Following cranial window implantation over dorsal CA1, animals were housed individually under standard conditions (12:12 h light/dark cycle, 22 °C, ad libitum food and water).

## METHOD DETAILS

### Surgery

Surgical methods are reported in Inayat et al. (2023)^2^. In brief, mice were anesthetized with isoflurane, and a 3 mm craniotomy was made over right dorsal CA1 (−2.0 mm AP, +1.8 mm ML) and overlying cortex was aspirated. A glass-bottom steel cylinder was implanted and sealed with dental acrylic, and a titanium head-plate was affixed for head fixation. Animals recovered for 3-8 weeks before behavioral training and subsequent testing with Brake/No-Brake behavioral paradigm.

### Behavioral Paradigm

Behavioral methods and training have been previously reported in Inayat et al. (2023)^2^. Briefly, animals were gradually habituated to head fixation over three days, with two sessions per day increasing from 5 to 30 min. Following habituation, mice were trained to run on a 150 cm linear, non-motorized Velcro conveyor belt (Country Brook) while head-fixed. The belt was blank and cleaned after each session, and all training occurred in darkness. Mice were trained daily (20-30 min) to run in response to a continuous, mild air stream directed at the back. The air was delivered via a solenoid valve from a regulated compressed-air source and terminated once the mouse ran a preset distance. After a 15 s inter-trial interval, the next air stream began. The required running distance was increased gradually from 30 cm to 150 cm (one full belt rotation), contingent on performance (>7 cm/s average speed), until mice achieved criterion performance (>75% successful trials) within three sessions. After training, mice were re-tested 10-18 days later, then performed a seven-configuration sensorimotor paradigm during two-photon calcium imaging. In the No-Brake (locomotion-permitted) condition, the air stream remained on until the mouse ran a fixed distance of 150 cm (air-on phase), followed by a 15 s air-off phase. In the Brake (immobility) condition, the air stream lasted 5 s (air-on phase), followed by a 10 s air-off phase. Behavioral signals including belt rotation (rotary encoders), lap reference (photo-sensor), and air, Brake, and light event timings, were synchronized with imaging data using a Digidata 1322A system (Molecular Devices Inc) and Arduino controller.

### Two-Photon Calcium Imaging

Detailed imaging procedures have been described in (Inayat et al., 2023). Briefly, CA1 activity was recorded using a Bergamo II multiphoton microscope (Thorlabs) with Ti:Sapphire excitation at 920 nm and a 16×/0.8 NA objective. Imaging was performed at 29.16 Hz (single-plane) or 9.72 Hz (dual-plane) with a 418 × 418 µm field of view. Laser power at the sample was ∼100 mW.

### Data Analysis

As reported previously in Inayat et al. (2023)^2^, images were registered and regions of interest (ROIs) corresponding to individual cell bodies were detected using Suite2P^65^; http://mouseland.github.io/suite2p. The number of cells identified with Suite2P from 5 animals were 341, 315, 315, 338, and 259 (total: 1568 cells) while they experienced the seven behavioral configurations. Cell percentages are reported with respect to the total number of cells identified with Suite2P.

Raw calcium traces for each ROI were obtained by averaging the fluorescence signal across all pixels within the ROI, followed by correction for neuropil contamination. For each fluorescence trace (*F*), a baseline value (*F*) was estimated, and the relative fluorescence change (Δ*F*/*F*) was computed as (*F* – *F*)/*F* . The resulting Δ*F*/*F* traces were then deconvolved to estimate neuronal firing rates (FR) using a first-order autoregressive model combined with constrained nonnegative matrix factorization^67^. All subsequent analyses of deconvolved activity traces were performed using custom MATLAB scripts (MathWorks).

The FR raw time series were binned using 0.3 s time bins and 3 cm distance bins. Time and speed modulation of FR was determined from time-binned data and distance modulation of FR was determined from distance-binned data. FR traces within a phase (air-on or air-off) were concatenated across trials before computing Pearson correlation and mutual information metrics (see below).

However, for visualization and calculating response fidelity (RF), raster plots were generated by plotting FR of individual trials versus time (time-bins) or distance (distance-bins). RF was defined as the percentage of trials (in the time-binned data) in which a neuron’s FR exceeded zero in at least one bin.

Peri-event time histograms (PETHs) were obtained by averaging FR across trials. Rate vector maps were generated by plotting the PETHs of all recorded neurons after normalizing each cell’s response to its peak firing rate. To assess population-vector level similarity, population vector correlations were computed by calculating the Pearson correlation between every pair of columns in the rate vector map. Each pixel in the resulting matrix represents the correlation between two population activity patterns, producing a symmetric matrix across the diagonal.

To quantify modulation by time, distance, or speed, we computed both Pearson correlation (PC) and mutual information (MI) between FR (y) and the variable of interest (x). For PC, we used the Pearson correlation of y with x. For MI, x and y were discretized into 10 bins and MI was computed on the joint distribution. Statistical significance for both metrics was assessed via permutation testing with 1,000 shuffles of FR (x held fixed), yielding a null distribution from which z-scores (zPC, zMI) and p-values were derived. Cells were deemed significantly modulated when the metric’s z-score exceeded the 95th percentile of the shuffle distribution (one-tailed, p < 0.05 or zPC/zMI > 1.645). All analyses were implemented in MATLAB using custom code.

### Single-Cell Identity Dynamics

To assess the stability and organization of modulation identities across configurations and air phases, we quantified overlap between cell modulation patterns using the Jaccard Index (JI). Each neuron was assigned an identity vector representing its modulation profile for time (T), distance (D), and speed (S). For each of the six cases in the No-Brake condition (three configurations × two air phases), the identity vector for each cell was defined as a binary triplet [T,D,S] where each element was set to 1 if the neuron was significantly modulated for that variable (based on permutation-tested Pearson correlation or mutual information metrics) and 0 otherwise. Similarly, for the Brake condition, a binary doublet [T,M] was defined for four cases (two configurations x two air phases). Pairwise similarity of identity vectors across the six cases was computed using the JI:

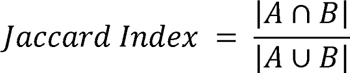

where *A* and *B* denote the binary identity vectors for the same cell in two different cases. For each cell, all 15 unique pairwise Jaccard values were computed and averaged to produce a mean agreement score. JI values can range between 0 and 1 with 1 indicating complete match. To test for phase-specific organization, we calculated the difference between average within-phase (air-on ↔ air-on, air-off ↔ air-off) and cross-phase (air-on ↔ air-off) JI values (Δ = same-phase − cross-phase). Statistical significance of Δ was assessed across animals using a one-sided Wilcoxon signed-rank test (α = 0.05).

### Population-level Identity Dynamics

To examine the organization of modulation identities at the population level, we quantified the overlap between groups of singularly modulated neurons (time-, distance-, or speed-modulated) across configurations and air phases using the JI. For each animal, population vectors were constructed for the 18 cases in the No-Brake condition representing the combination of three modulation types (T, D, S), three configurations (C3, C4, C5), and two air phases (air-on, air-off) as well as 8 cases for the Brake condition with two modulation types (T, M), two configurations (C2, C7), and two air phases (air-on, air-off). Each population vector corresponded to the binary identity of cells belonging to a particular modulation type in a specific configuration and phase. Pairwise Jaccard indices were then computed between all possible pairs of population vectors using the same relation as above but with A and B denoting the sets of cells identified as modulated in the two cases being compared. This yielded an 18 × 18 population agreement matrix for each animal for the No-Brake condition, in which each cell of the matrix represented the similarity between two modulation-defined populations. When considering both Brake and No-Brake conditions, 26 × 26 population agreement matrix was generated for each animal. Matrices were averaged across animals to obtain group-level representations.

To assess higher-order structure, the Jaccard distance (1 − JI) was used as a dissimilarity metric for agglomerative hierarchical clustering (MATLAB linkage function, unweighted average linkage). sThis approach grouped together populations with similar modulation profiles across configurations and air phases. Cluster quality was evaluated using the cophenetic correlation coefficient (c), which measures how well the hierarchical tree preserved pairwise distances among populations.

## QUANTIFICATION AND STATISTICAL ANALYSIS

Statistical analysis was done in MATLAB® 2024a. Data are reported as mean ± standard error of mean (SEM). Unless otherwise noted, repeated-measures ANOVA (RM-ANOVA) was used to compare parameters across conditions/cases (n = 5 animals) with various within-subjects factors as reported in the text. Normality was assessed with the Shapiro-Wilk test (“swtest.m,” MATLAB File Exchange), and sphericity with Mauchly’s test; Greenhouse-Geisser corrections were applied where violated. Minor deviations from normality and isolated outliers were tolerated given the within-subject design. Post hoc pairwise comparisons were conducted using Tukey’s HSD (α = 0.05). Significance levels are denoted as p < 0.05 (*), p < 0.01 (**), and p < 0.001 (***). Effect sizes are reported as partial η^2^ = (Sum of Squares _effect_)/(Sum of Squares _effect_ + Sum of Squares _error_) to complement p-values given the small sample size (n = 5).

## AUTHOR CONTRIBUTIONS

All authors participated in the design of this study, with S.I. and I.Q.W. as the main contributors. S.I. and B.B.M. performed the experiments. S.I. conceived and developed the air-induced running task. S.I. performed the data analyses with participation of the other authors. S.I. and I.Q.W. wrote the manuscript, which all authors commented on and edited. M.H.M. supervised the study.

## ACKNOWLEDGEMENTS

We thank Dr. JianJun Sun for performing animal surgeries, Di Shao and the University of Lethbridge Animal Care Services staff for animal husbandry. We thank Adam Neumann and Dr. Maurice Needham for technical assistance with two-photon microscopy. This study was supported by the following research grants to Dr. Majid Mohajerani: Canadian Institutes of Health Research (390930), Natural Sciences and Engineering Research Council of Canada (40352), Alberta Innovates (43568), Alberta Alzheimer Research Program Grant (PAZ15010), Alberta Alzheimer Research Program Grant (PAZ17010), Alzheimer Society of Canada (43674). The funders had no role in the study design, data collection and interpretation, or the decision to submit the work for publication.

